# The synergy between cannabidiol and probiotics curtails the glycemic indicators, enhances insulin production, and alleviates the symptoms of type 2 diabetes

**DOI:** 10.1101/2024.06.04.597375

**Authors:** Sahar Emami Naeini, Bidhan Bhandari, Hannah M Rogers, Jules Gouron, Pablo Shimaoka Chagas, Lívia Maria Maciel, Henrique Izumi Shimaoka Chagas, Jack C Yu, Mohammad Seyyedi, Évila Lopes Salles, Mark Fields, Babak Baban, Lei P Wang

**Author notes:** **Corresponding authors:** Lei P Wang, Ph.D., -Department of Oral Biology and Diagnostic Sciences, DCG., -DCG Center for Excellence in Research, Scholarship and Innovation (CERSI) Augusta University, Augusta, GA, USA., Babak Baban, Ph.D., MPH, MBA, FAHA., -Department of Oral Biology and Diagnostic Sciences, DCG., -DCG Center for Excellence in Research, Scholarship and Innovation (CERSI) Augusta University, Augusta, GA, USA.

## Abstract

Diabetes continues to challenge healthcare system as one of the most growing epidemics with staggering economic burden. It is estimated that 783 million by 2045 will live with diabetes worldwide, 90% of those cases are type 2 diabetes (T2D). T2D is a multifaceted disease, its treatment requires a holistic approach, beyond single target medications with high efficacy. There is a dire need to explore and invent new and effective therapeutic modalities for T2D.

In this study we tested whether a combined formulation of cannabidiol (CBD) and probiotics could control glycemic indices and alleviate symptoms of T2D. We used a mouse model of T2D, replaced their drinking water with a combination of CBD and probiotics formulated as a commercially available beverage.

Our findings demonstrated that combination of CBD and probiotics not only reduced the glycemic indices (HbA1c & FBG), but also altered the microbiome profile, promoted beneficial bacteria. Further, the CBD/probiotic combination reduced peripheral inflammatory cytokines and enhanced insulin production in pancreatic islets.

In conclusion, our results suggest that consumption of combined CBD and probiotics could be used as a natural, practical, affordable, and safe alternative and complementary therapeutic modality to treat T2D.

## Introduction

Type 2 diabetes (T2D), known as the “silent epidemic” is emerging as one of the most pressing health-related challenges of our time (1). Globally, there are over half a billion individuals living with diabetes, of those, 96% are T2D (2-3). Given the estimate of 1.3 billion diabetics by 2050 (4), there is a dire need for new therapeutic strategies to prevent and treat diabetes. Despite significant advances in the treatment of diabetes through medications, there are growing high demands by public and industry for novel and alternative diabetic modalities based on natural approaches rather than chemical compounds and medications. This is mainly because of potential side effects as well as many unknown about current medications and high impact of dietary habits on the progression of T2D.

Cannabidiol (CBD) is one of the major phytocannabinoids produced by cannabis plants. CBD is a non-psychoactive cannabinoid which makes it generally safe compound compared to tetrahydrocannabinol (THC), the other well-known cannabinoid with psychoactive effects. Recent work by our laboratory and others suggests beneficial effects of CBD alone or in combination with other cannabinoids and compounds in the treatment of a wide spectrum of pathologic conditions and diseases (5-6). Several studies have proposed that CBD could be beneficial in the treatment of diabetes (7-8) warranting further research.

Gut microbiome and its impact on metabolic diseases such as diabetes has been extensively investigated (9-10). Interventional strategies such as microbiome graft and dietary probiotics have shown beneficial effects in diabetes, suggesting a potential therapeutic connection between microbiome manipulation and improving the symptoms of diabetes.

In this study, we investigated if a combination of CBD and probiotics commercially available as a beverage (CANABIX) could alleviate the symptoms of T2D. To test, we used a mouse model of T2D in a preclinical setting. Our findings demonstrated that a combination of CBD and probiotics could improve the symptoms of T2D significantly, suggesting a highly novel therapeutic application for CBD in the treatment of T2D.

## Material and methods

Fourteen-week-old male db/db mice were obtained from the Jackson Laboratories. The db/db mice are well-established/accepted mouse models to study diabetes type II and obesity. The use of animals for these studies were in accordance with ethical standards and was approved by the Institutional Animal Care and Use Committee at Augusta University. Diabetic mice then were divided into two experimental groups. In one group, the daily drinking water was replaced by CANABIX, a beverage containing a combined CBD (30 mg) and probiotics (*Bacillus coagulans* MTCC 5856, LactoSpore®, Sabinsa USA) formulation, provided by Hydro One beverages, LLC (Greenwood SC, USA). The second group continued drinking regular daily water. The mice (n=8/group) were monitored for 9 weeks drinking between 45-50 ml per week.

### Glycemic measurements, microbiome assessment and bioassays

Fasting blood glucose (FBG) and hemoglobin A1c (HbA1c) were measured weekly by blood collection through tail. Fresh fecal samples (2-3 pellets/mouse) were collected, immediately frozen, and stored in a –80 °C freezer until analysis. DNA Extraction, Library Preparation, Sequencing DNA extraction and library preparation were performed by Novogene Lab (Durham, NC, USA). For flow cytometry analysis, fresh whole blood was collected, processed and intracellular staining of TNFα and IL-17 (BD BioSciences, Bedford, MA, USA) was performed. Samples were then run through a a NovoCyte Quanteun and analyzed by FlowJo analytical software. The FlowJo plugin for the algorithm “uniform manifold approximation and projection” (UMAP) was used to perform dimension reduction of down-sampled and concatenated data sets. Further, freshly harvested pancreas tissues from db/db mice were collected and processed for Immunohistochemistry staining and imaging by using bright field Zeiss (AXIO Imager M2) light microscope and ImageJ software (version 1.53e) for area intensity analysis. For statistical analysis, Brown-Forsythe and Welch analysis of variance (ANOVA) was used to establish significance (*P*<0.05) among groups. For tissue quantification statistical analysis, we compared the area of expression in both treated with beverage and non-treated groups by using two-way ANOVA followed by post-hoc Sidak test for multiple comparison (*P*<0.05).

## Results

### Combination of CBD and probiotic showed antidiabetic activities, reduced HbA1c and plasma glucose and altered microbiome profile in mice with T2D

Daily drinking of combined CBD and probiotics resulted in significant reduction of HbA1c (from 7% to 4%, *P*<0.05) and lower FBG (from 306 to 235, *P*>0.05) compared to drinking regular water (Figure 1A). Further, daily drinking of CBD and probiotics combination resulted in significant alteration of several bacterial phyla within gut microbiome. While changes were observed for most frequently phyla, however, Dubosiella and Lactobacillus showed the most significant increase compared to the control diabetic mice drinking regular water (Fig 1B-C).

**Fig 1.**
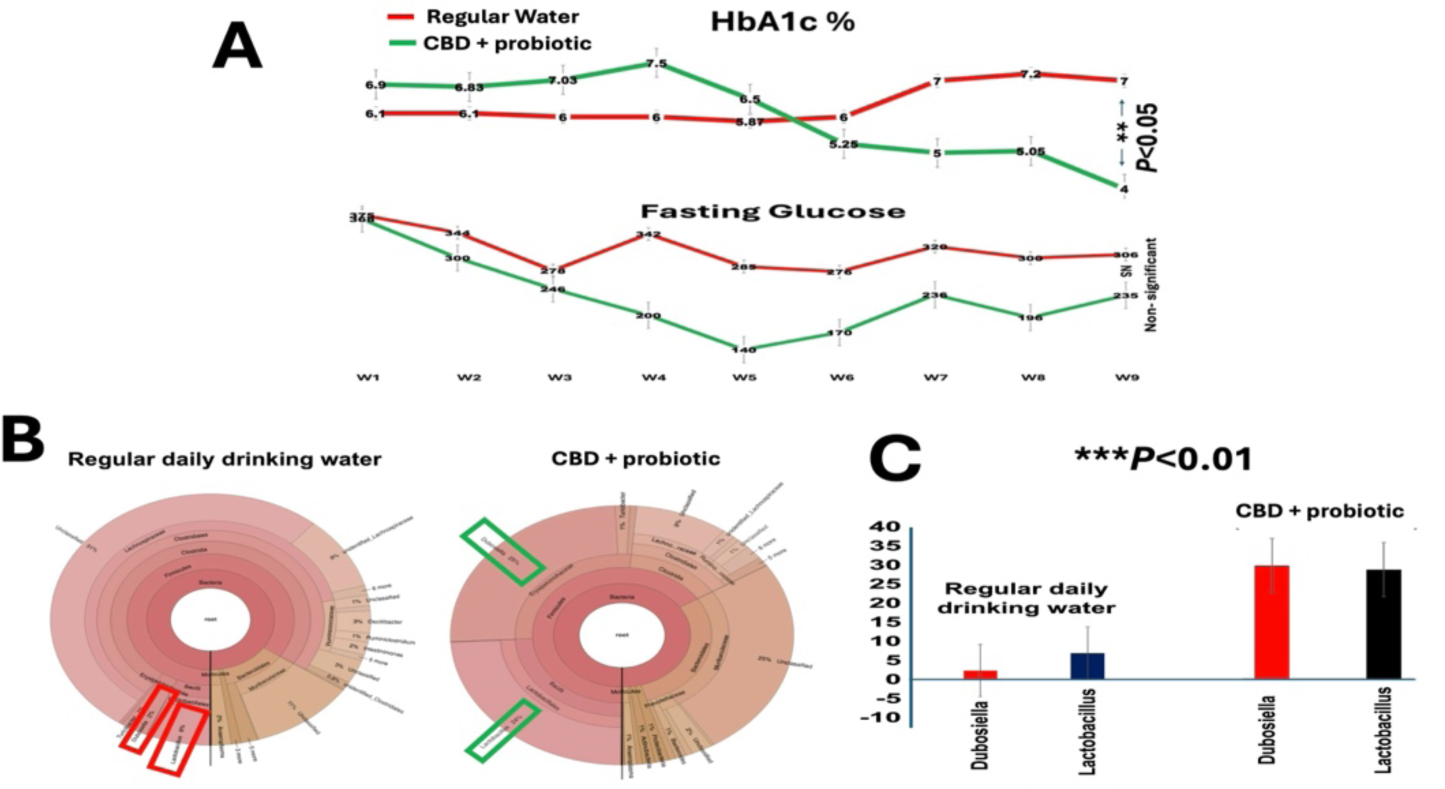
Combination of CBD and probiotic reduced glycemic indices and altered gut microbiome. **A)** Daily drinking of beverage containing CBD and probiotics resulted in significant reduction of HbA1c (from 7% to 4%, P<0.05) and lower FBG (from 306 to 235, P>0.05) compared to drinking regular water. **B)** Consumption of combined CBD and probiotic formulation resulted in significant alteration of several bacterial phyla within gut microbiome. While changes were observed for most frequently phyla, however, C) Dubosiella and Lactobacillus showed the most significant increase compared to the control diabetic mice drinking regular water (*P*<0.01 for both bacteria).

### Combination of CBD and probiotic regulated proinflammatory responses and enhanced insulin production in mice with T2D

Daily drinking of CANABIX beverage resulted in significant reduction of proinflammatory cytokines, TNFα and IL-17 (Fig 2A-B) improving insulin resistance (11-12). The flow cytometry UMAP (Fig 2A) displays numerous spatial clusters with significant differences in distribution between diabetic mice (db/db) drinking CBD/probiotic combination versus mice drinking regular daily water only. Bargraphs demonstrates the quantified measures for TNFα and IL-17. Further, drinking CBD and probiotic combination resulted in significant increased in the insulin production (*P*<0.01) in the pancreatic islets of diabetic mice (Fig 2C-D).

**Fig 2.**
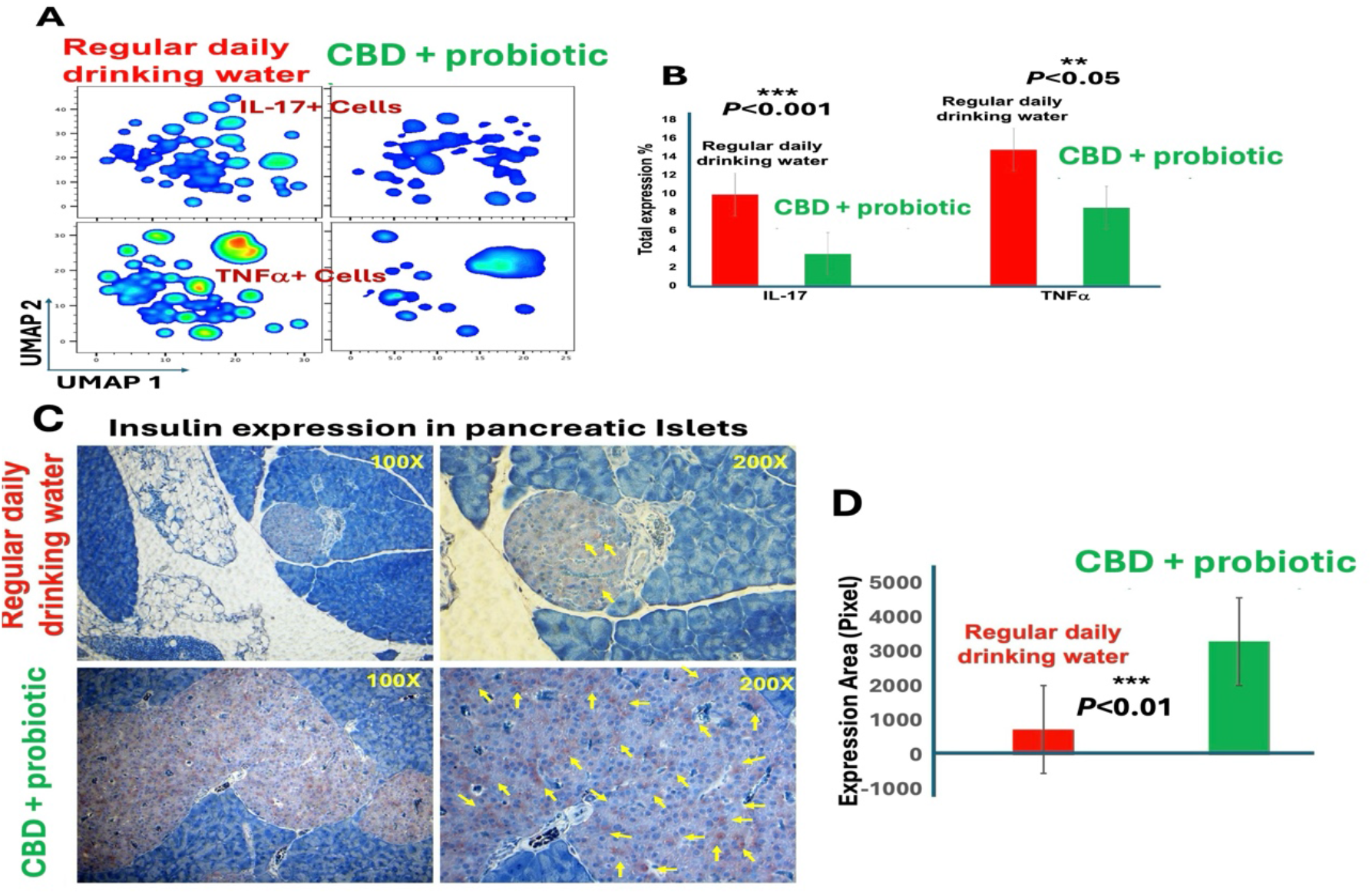
Combination of CBD and probiotic diminished proinflammatory responses and enhanced insulin production in diabetic mice. **A)** Flow cytometry UMAP revealed numerous spatial clusters with significant differences in distribution of proinflammatory cytokines including TNFa and IL-17, between mice with T2D consumed combined CBD and probiotic versus mice used regular daily water only. **B)** Bargraphs demonstrates the quantified measures for TNFa and IL-17 (P<0.05). **C)** lmmunohistochemistry staining for insulin in diabetic mice consumed CBD and probiotic showed significant increase in the insulin production (P<0.01) in pancreatic islets of diabetic mice used daily CBD and probiotic (Yellow arrows) versus diabetic mice using drinking water only. **D)** Quantified measures of insulin production displayed a significant difference (P<0.01) between insulin production in the pancreatic islets of diabetic mice consumed daily CBD and probiotic versus diabetic mice using drinking water only.

## Discussion

The novel findings here in this current study are the first to demonstrate a significant beneficial effect of a combination of CBD and probiotics in a preclinical model of T2D. A significant reduction of HbA1c and lower FBG in a period of 6 weeks through a non-invasive supplementary beverage is a breakthrough and a potential game changer in the field of diabetic care.

Diabetes continues to threaten global health, remains as a major economic burden, incapacitates working individuals worldwide, affecting life quality and causes mortality (1). During last two decades, world has witnessed major advancements in the treatment of T2D including introduction of insulin analogs, development of new ways of insulin delivery systems, pharmacologic innovations (e.g., GLP-1 agonists etc.), genetic approaches and even psychologic and behavioral therapies (13-14). Despite all achievements, diabetes is expected to rise to a towering level of 643 million by 2030 and the total number of 783 million by 2045 worldwide (1,4), 90% of those will be T2D. Such exponential growth is simply because T2D is a disease with multifaceted nature and several factors including diet, dehydration, medications, stress, sicknesses, pain, physical activities, general lifestyle and importantly, the socioeconomic status will influence the T2D management and clinical outcomes (15-16). Given the high cost and time of developing a drug, sideeffects, issues with availability and monitoring, it is thus a dire need to explore and invent alternative strategies to prevent and treat T2D in affordable, safe, and more effective ways.

The beneficial effects of CBD combined with probiotics observed during this study were through consuming a dietary beverage supplement. Considering the central role of hydration in alleviating symptoms of T2D (17), such delivery strategy by beverage magnifies the significance of novel findings in this study. Further, the probiotic content of the beverage included culture of beneficial bacteria shown to improve prediabetes symptoms and to help digestion. Among all bacteria, two species of Dubosiella and Lactobacilli demonstrated a significant increase in diabetic mice used CANABIX beverage compared to the diabetic mice consumed regular drinking water. Dubosiella is reported to have anti-aging functions by reducing oxidative stress and to improve vascular endothelial function (18). Lactobacilli are known to have inhibitory effects on harmful bacteria by producing lactic acid, significantly reduce HbA1c, controlling fasting insulin levels, and regulating HOMA-IR levels in T2D (19).

Importantly, individuals with T2D are at higher risk of anxiety (20). CBD is known as a potent regulator of inflammation and an effective anxiolytic agent which could help people with T2D to maintain and continue their daily life with higher quality.

Further, our findings showed that consumption CBD and probiotics combination induced a significant reduction in the expression and distribution patterns of IL-17 and TNFα. Both IL-17 and TNFα are prominent inflammatory cytokines, well known for their contribution and roles in the development of insulin resistance (11-12). Therefore, downregulation of both IL-17 and TNFα by a beverage could suggest a very effective, practical and affordable strategy to improve and control the glucose level in diabetes. More importantly, using combination of CBD and probiotics increased insulin production in pancreatic islets of db/db mice. This is highly significant discovery since it can effectively control the blood glucose and improve the clinical outcomes profoundly. This augmentation of insulin production by a beverage is remarkable due to the lack of efficacy, inadequacy and cost of current treatments in boosting insulin production effectively in diabetic patients.

In conclusion, the beneficial effects observed in this study present an alternative, practical and novel therapeutic modality in the treatment of T2D with minimal to zero side effects, in a non-invasive and affordable fashion. Obviously, there are limitations in current study such as power analysis and experimental groups, warrant further research and justifies a human clinical trial.

## Abbreviations

CBD: Cannabidiol
FBG: Fasting Blood Glucose
GLP-1: Glucagon-like peptide 1
HbA1c: Hemoglobin-A1c
HOMA-IR: Homeostasis model assessment for insulin resistance
IL-17: Interleukin-17
THC: Tetrahydrocannabinol
TNFα: Tumor necrosis factor alpha
T2D: Type 2 Diabetes
UMAP: Uniform Manifold Approximation and Projection.

## Author approvals

All authors have seen and approved the manuscript, and that it hasn’t been accepted or published elsewhere.

## Acknowledgement

Authors are thankful to Hydro One Beverages (SC USA) for providing the CANABIX drink for this study. Authors also thank Medicinal Cannabis of Georgia for providing help in design and optimizing the CBD and probiotic dosage.

## Data availability statement

All data supporting the findings of this study are available within the paper and also are available upon request from the corresponding author.

## Ethical publication statement

We (authors) confirm that we have read the Journal’s position on issues involved in ethical publication and affirm that this report is consistent with those guidelines.

